# PBrowse: A web-based platform for real-time collaborative exploration of genomic data

**DOI:** 10.1101/068049

**Authors:** Peter S. Szot, Andrian Yang, Xin Wang, Uwe Röhm, Koon Ho Wong, Joshua W. K. Ho

## Abstract

**Summary:** The central task of a genome browser is to enable easy visual exploration of large genomic data to gain biological insight. Most existing genome browsers were designed for data exploration by individual users, while a few allow some limited forms of collaboration among multiple users, such as file sharing and wiki-style collaborative editing of gene annotations. Our work’s premise is that allowing sharing of genome browser views instantaneously in real-time enables the exchange of ideas and insight in a collaborative project, thus harnessing the wisdom of the crowd. PBrowse is a parallel-access real-time collaborative web-based genome browser that provides both an integrated, real-time collaborative platform and a comprehensive file sharing system. PBrowse also allows real-time track comment and has integrated group chat to facilitate interactive discussion among multiple users. Through the Distributed Annotation Server protocol, PBrowse can easily access a wide range of publicly available genomic data, such as the ENCODE data sets. We argue that PBrowse, with the re-designed user management, data management and novel collaborative layer based on Biodalliance, represents a paradigm shift from seeing genome browser merely as a tool of data visualisation to a tool that enables real-time human-human interaction and knowledge exchange in a collaborative setting.

**Availability:** PBrowse is available at http://pbrowse.victorchang.edu.au, and its source code is available via the open source BSD 3 license at http://github.com/VCCRI/PBrowse.

**Supplementary Information:** Supplementary video demonstrating collaborative feature of pbrowse is available in https://www.youtube.com/watch?v=ROvKXZoXiIc.

## 1 INTRODUCTION

Consider this situation: A group of scientists from various institutions around the world participates in a project that involves generating new genome-wide ChIP-seq, RNA-seq and whole genome sequencing data. The scientists involved in data generation and data analysis are located in different countries. At regular intervals, meetings are held to discuss the data and share analysis results. To help visualise the data, the bioinformaticians need to take screenshots of the genome browser view at many different resolutions and email them to their colleagues. Even some browsers such as Ensembl and UCSC support persistent views that can be shared by URL in the later update, it is stilla time consuming and repetitive task. Also, in this situation, the responsibility of the analysis is entirely placed on the shoulder of the bioinformatician, who may not be able to fully utilising the expertise available in the collaborative research group.

Many of the current genome browsers were developed to facilitate individualised exploration of genome-scale data. Nonetheless, as illustrated in the above example, an individualised exploration of data may not be fully harnessing the wisdom of the crowd within a research consortium. As this type of collaborative projects is becoming more commonplace, it is increasingly important to facilitate real-time collaborative exploration of genome-scale data. The central goal of this work is to develop an intuitive web-based platform where genome-scale data can be shared and visualised collaboratively in real-time. In this paper, we discuss how we implement a real-time collaborative genome browser that fulfils this need.

### Evolution of the genome browser technology

Genome browsers were developed out of a need to simplify the visualisation and analysis of an increasingly vast amount of genomic data, and make them accessible for all research communities to share and expand upon. The first generation of genome browsers followed the dynamic-server static-client model of content delivery, as illustrated in Figure 1A, working on the assumption that the client is likely a slow machine, incapable of performing complex operations. Thus, the server would necessarily prepare and pre-render all elements of a page before delivering it to the client, which would then simply display it as static content. The human genome browser at the University of California at Santa Cruz (UCSC) was one of the first browsers implementing this model (1). It allows for a reliable display of any portion of the genome, at any scale. It also enables a simple form of collaboration by allowing the upload of annotation tracks to be viewed by other research groups. While it was originally named as a “human” genome browser, it has over the years been upgraded to enable visualisation of data from various species. The Ensembl genome browser was another major system (2). The intention of the Ensembl project was to offer an integrated, extensible, and reusable framework for generating, storing, retrieving, and displaying genomic annotation data. Same as UCSC genome browser, it has since received a multitude of updates and has grown dramatically G-Compass is another browser following this client-server model (3) but attempts to address more specific problems than either UCSC Genome Browser or Ensembl. The developers sought a way to resolve the problem of comparing multiple disparate genomes simultaneously while in a meaningful way. The goal of their browser is to provide effective comparisons between human and selected model organisms, promoting the exchange of functional information between different organisms. With the advent of better client-sided technologies for retrieving and processing data through in-browser Java scripting, the client is no longer completely reliant on the web-server to pre-process data. This effectively frees the server from a significant burden of work, while utilizing the enhanced power of the client computers to achieve greater responsiveness, and reduce the server overhead, as shown in Figure 1B. JBrowse is one such genome-browser (4). Utilising the computational power of the client side machine, JBrowse enables smoothly animated panning, zooming, navigation, and track selection. This important feature preserves the user’s sense of location by avoiding discontinuous transitions.

**Figure 1.**
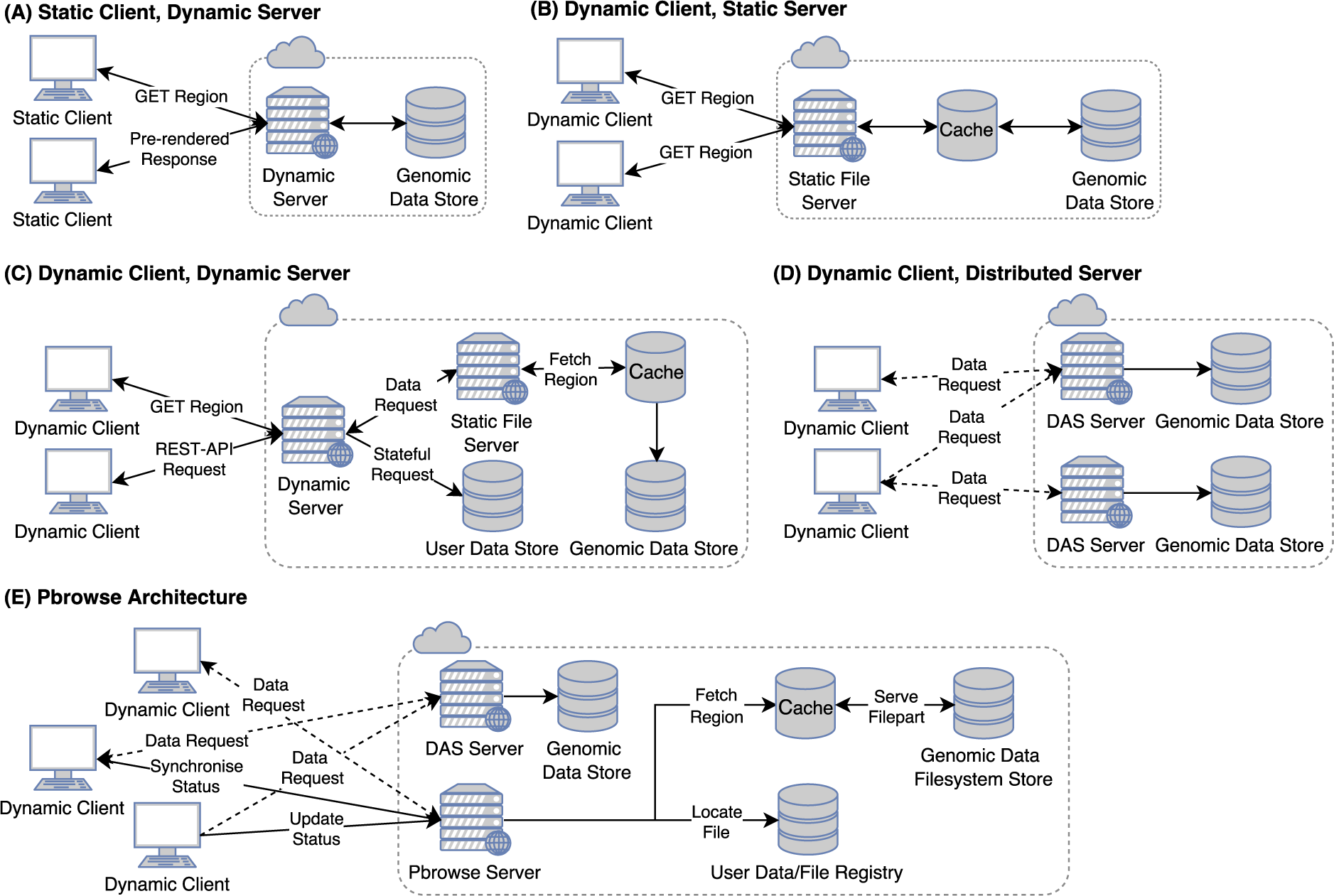
Comparison of different system architecture of web-based genome browsers.

While a dynamic client is capable of performing a great deal of processing, it is limited in what it can achieve alone. Adding a dynamic server allows for the implementation of significantly more complex functionalities and persistent cross-session interactions between multiple clients (Figure 1C). Developers of ABrowse (5) adopted this model, extending the efficient browser framework established by JBrowse, by enhancing interactivity, opening access to more data sources, and providing support for inter-user collaboration in terms of enabling non-real-time commenting and annotation. Genome Maps (6) is another such system, designed specifically to address the browser efficiency issues under increasing data loads brought about by high-throughput sequencing technology. It aims to provide a near seamless browsing experience through real-time navigation along chromosomes, all the way down to individual base pairs. GBrowse, similar to Genome Maps, was designed specifically to handle the large scale of Next Generation Sequencing (NGS) datasets (7). ChromoZoom is yet another browser that handles high volume NGS data with a focus on using modern technologies to improve the user experience. It facilitates the exploration of experimental data by researchers, enabling the visualization of custom results alongside a dynamic representation of curated genomic information (8).

With the ever increasing power of client-sided rendering technologies, the role of the web-server as a central point for accessing data is becoming increasingly redundant. As data sets continue to grow, storage of genomic data on a single server becomes problematic. One solution is to distribute the load. An example of this is depicted in Fig 1D. The Dalliance browser (9) addresses these concerns, opting for a completely self-contained and embeddable module approach, offering high levels of interactivity which is competitive with specialized desktop applications such as the Integrative Genomics Viewer (IGV) (10), all the while running entirely within the web browser.

### Design of a real-time collaborative genome browser

To date, most genome browsers focus on exploration of data by individual users. To harness the wisdom of the crowd, we propose to introduce real-time collaborative technology into the genome browser itself, in order to streamline the process of discovery, exploration, and the subsequent sharing of results, by eliminating the need for third party software solutions.

With regards to real-time collaboration, many genome browsers mentioned previously have some elements of collaboration integrated. However, they are always limited in some regards. In some systems it may be possible to share a view via URL or upload an annotation track for other users to make use of; none of which can be considered as ‘real-time’ collaboration.

The concept of distributed real-time editing has existed in some forms for many years and numerous algorithms have since been proposed as solutions to the three primary problems: causality preservation, user intention preservation, and convergence. In the case of a collaborative genome browser, we assume the need of users to edit the raw data is limited. Instead, only the viewpoint and superficial changes to tracks are allowed. This reduces the complexity of the operation space which we must consider; in particular, insertion or deletion will have no effect and can be totally ignored. In order to address the requirement of high-responsiveness given non-deterministic latency, Sun et al. (11) have proposed a multi-versioning approach. Multiple conflicting operations are made permissible by allowing the creation of branching ‘copies’. This preserves the intention of all participating collaborators while providing optional consistency. In a browser context, shifting views while collaborating in different directions is a conflicting operation which will produce two diverged states. Each user will see the position of the others and can resynchronise themselves at will.

Another alternative is to utilise the idea of user-roles and access rights (12). In this way, the session owner is considered to have administrator privileges and is able to grant permissions to other participants, allowing or disallowing them to perform specific operations. This can prevent conflicts from ever happening if users are only permitted to perform compatible operations concurrently. In the browser context however, there are too few valid operations to requiresuch restriction. Instead the session leader is assigned the privilege to manage the session. By default, any updates made by the leader will override those made by followers, though this can be overridden by the follower who can choose to subscribe to the updates to any set of users, or none.

Finally, we observe a recent development of a collaborative web-based Java IDE (CoRED) (13) utilising many of the aforementioned techniques. They implement differential synchronization whereby the server stores the shared document, and each client has a separate shadow copy of the document both on server and client side, along with the copy they are editing. When edits are made, the changes are processed into a patch which is merged and redistributed by the server. Patches are small as they contain only the modified data, keeping a high responsiveness as well as enabling high concurrency. CoRED as a whole, serves as a useful example of the technologies utilized in a modern collaborative editor, many features of which will be essential to the success of the proposed project. PBrowse is an amalgamation of the best features of many widely used genome browsers and the latest technology for real-time collaboration, with the architecture as depicted in Figure 1E. It effectively addresses all the weaknesses of previous genome browsers, combining highly responsive and flexible collaborative features, with a completely transition free, and intuitive browsing experience. Sharing and exploring data collaboratively has been streamlined so that researchers can spend more time finding new insights.

## 2 MATERIAL AND METHODS

### System architecture

Since PBrowse was formulated with the idea of enabling real-time collaboration between distinct users, it necessitates their identification by user created accounts. A user can quickly and easily register for an account via the user interface of the web-application. A new user must enter a unique username and provide a valid email address – which is used for account verification and several other features. When logged in as a user, all actions are performed in the context of that user, with all their associated privileges.

The client communicates directly with a primary server, the synchronisation point between all active clients. Where PBrowse differs from other genome browsers, is in its use of the WebSocket as a communications channel. The browser is served over a plain HTTP connection from a servlet endpoint, important for the restriction of most web-browsers in loading mixed content, i.e. HTTP, over HTTPS. PBrowse hosted data is securely accessible, but the majority of genomic data providers serve data only over HTTP. To run PBrowse over a secure channel would limit the availability of a large portion of these data. Instead, only the WebSocket connection is necessarily run over an encrypted protocol, ensuring the confidentiality of user submitted information such as email addresses and passwords.

PBrowse depends on a JSON-encoded message-based API with the client and server both recognising a discrete set of strings which each trigger a unique function. Most of the time, messages are sent to the server when the client performs either a synchronised action, or requests state information. The message flow between client and server is depicted in Figure 2. The server responds with a predefined message type which the client asynchronously processes upon its reception. Some client actions require updates to be propagated to multiple users, the WebSocket allows the server to notify these users in a “push” style, without the need for a client to request updated data. This both reduces the bandwidth overhead of synchronisation, and greatly reduces the latency of updates. The client alone recognises over 35 different message response types, with several messages initiating multiple different functions depending on the given parameters.

**Figure 2.**
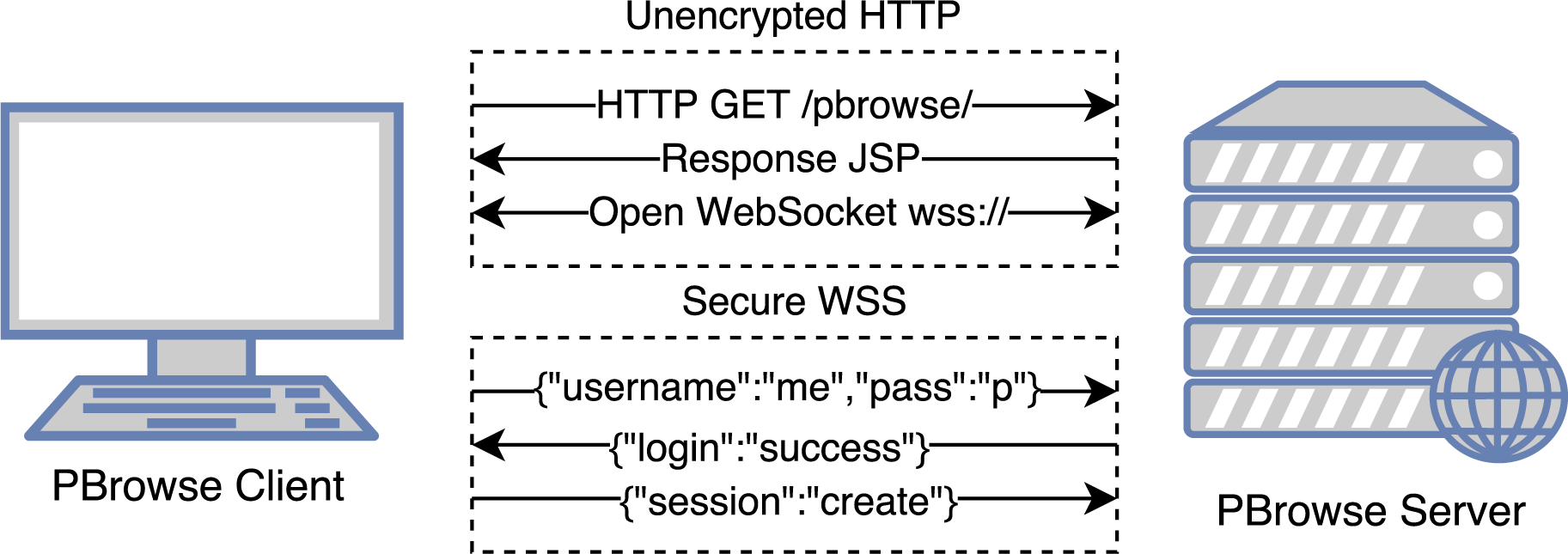
User login workflow. Transition from HTTP transfers to WebSocket SSL communication.

The login request is one such message, which if successful, associates the WebSocket connection of the calling user, with the authenticated account information. The server also generates a unique token which is passed to the client and stored as a cookie. Whenever the client’s connection is interrupted – for whatever reason, it will attempt to perform a session resume by transmitting the cookie to the server. If the token information matches the server’s copy, the calling user is automatically logged in, otherwise the client will discard the data and manual login is required once more. Similarly, only one user may be logged into a particular account at any one time. In the event of multiple logins, the last user to successfully authenticate is granted access to the account; all others are automatically logged out. If the user forgets her/his password, she/hemay request a reset which only requires access to the email address used in account registration.

### File storage and database

PBrowse makes use of a MySQL database instance, used not to store track data, but rather only meta-information. For instance, when a user uploads a new track to PBrowse, the user-provided metadata is saved as an entry in the database – a Data Descriptor (DD). The file itself is saved to disk in a predetermined location. The path to the file is saved as part of the DD for the uploaded file and recalled whenever the data is requested.

All other persistent data storage is managed entirely through the database. The user account, and more specifically, the username, is the key which connects a particular user to all of the files she or heowns, the groups she/he is member of, and the comments she or he has written. Requests to access any of these data occur via a set of parameterised SQL queries. The returned data is filtered depending on the requirement and is typically transmitted to the client as part of a JSON encoded message via the WebSocket.

When accessing genomic data for display in the genome browser, only the ID of the track must be known. The PBrowse server implements an REST-based endpoint for serving files, taking a single parameter – the fileID. For sources which require an index, adding an .ext to the request will cause the server to read the file with the path of the original request, appended with the extension. It is a convenient shortcut when the ID of the index file is not known.

First, however, the identity of the calling user must be verified. This is done using the user-uauth cookie pair which is transmitted by the user along with the file request. If it matches the pair stored by the server, their identity can be confirmed. The file server then requests the DD of the file and checks if the file is publicaly visible, or if the caller is the owner. If neither, it checks if the caller is in a collaborative session with the owner. Finally, it checks if the caller is part of a group with access to that file. If the caller is a member with the appropriate permissions, access is granted and the server fulfils the request for a specific range of data. Otherwise, if all of the checks fail, the server responds with a forbidden error.

Since the genome browser typically makes many requests for small ranges of data, there can be significant overhead when continuously retrieving DDs or grouping information. This was overcome via the use of caching. When a file with ID x is requested, the server first tries to recall it from the cache, using x as the key. If it is not found, the server will then request it from the database. The same is done for user group requests, mapped to the caller’s username. The cache entries never expire, to ensure that repeated reading will not necessarily waste CPU time. In the event that the file DD or group listing is modified after being cached, the cached entry is immediately invalidated, causing the next request to recall the information from the database again.

### Groups and file access control

The user group system is an add-on to the functionality of collaborative file sharing, with several more fine grain access control systems. Essentially, groups allow users to share data only with specific users, without the necessity of making the data public for all.

Any logged-in user is able to make a new group providing just a unique group name. Upon its creation, they are appointed the role of owner – which is the most authoritative role, capable of performing any group administration task. Aside from the ownership role, a user may be assigned any combination of the following privileges within the context of the group: file reading, file management, and user management. The owner automatically has all of these privileges and cannot lose them. If the owner of a group decides to leave it, the entire group is deleted. The group management interface is depicted in Figure 3.

**Figure 3.**
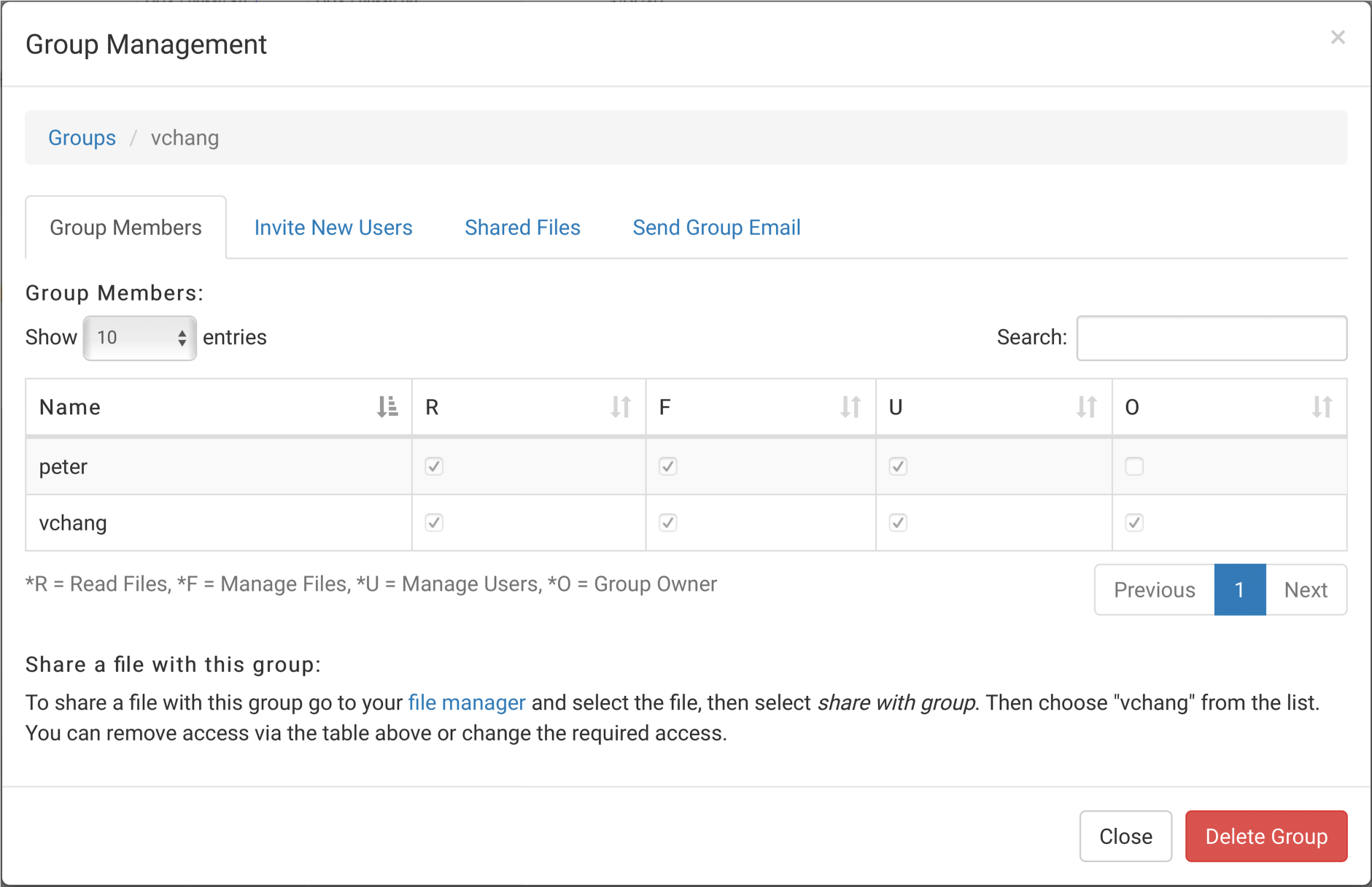
Group management interface. Allows for management of group, such as invitation or removal of users, modification to user permissions, access to shared files, and sending group message.

New users may be invited to join the group by any members currently having the user management privilege. If the invited user is currently logged in, they will receive notification of the invitation and will be granted immediate access to all of the group’s resources, depending on their privilege level. Otherwise, the user will see the new group in their group listing when next they log on.

In addition to inviting new users, user managers may also modify the permissions of other users in the group. There are three restrictions on this however: firstly, a manager cannot modify the user manager permission. Secondly, a manager cannot modify their own permissions, and thirdly, a manager cannot modify the group owner’s permissions. Only the owner can bypass restrictions one and two.

File managers, on the other hand, are only able to control group file access. These users are able to share their privately accessible files with the group. All group members with the file read permission are then able to view the data in the genome browser and leave comments on it. A single file may be shared with any number of groups, provided the file owner has the file management permission in each of them. A manager may also remove group file access, but only for files they have shared. The group owner can bypass this restriction and may remove any file from group access.

The file read permission, as the name implies allows users to access files shared with the group. These users are not able to perform any level of group administration. Note that it is possible for a user to have file management permission and not have file read permission. While unusual, any combination of the three permissions can be set, including no permissions which will allow group members only to see who else is in the group.

One final owner-only privilege is the send-group-message permission. This allows the owner to send out email messages to all members of the group, without ever revealing the users’ email addresses. This feature should only be used for high importance notifications or perhaps to arrange further collaborative sessions.

## 3 RESULTS

### The PBrowse genome browser

The front-end of PBrowse is based on the Dalliance genome browser (4). It is embedded and fully contained within a single webpage, with all browsing actions performable without the need for any extra navigation or refreshing of the view. The interface is designed with responsiveness in mind, allowing simultaneous and asynchronous loading of multiple data sources, to ensure transitions when viewing different parts of the genome are as seamless as possible. Data for regions upstream or downstream of the viewed genomic region are loaded in advance in anticipation of a user panning the view. An example of the collaborative view can be seen in Figure 4.

**Figure 4.**
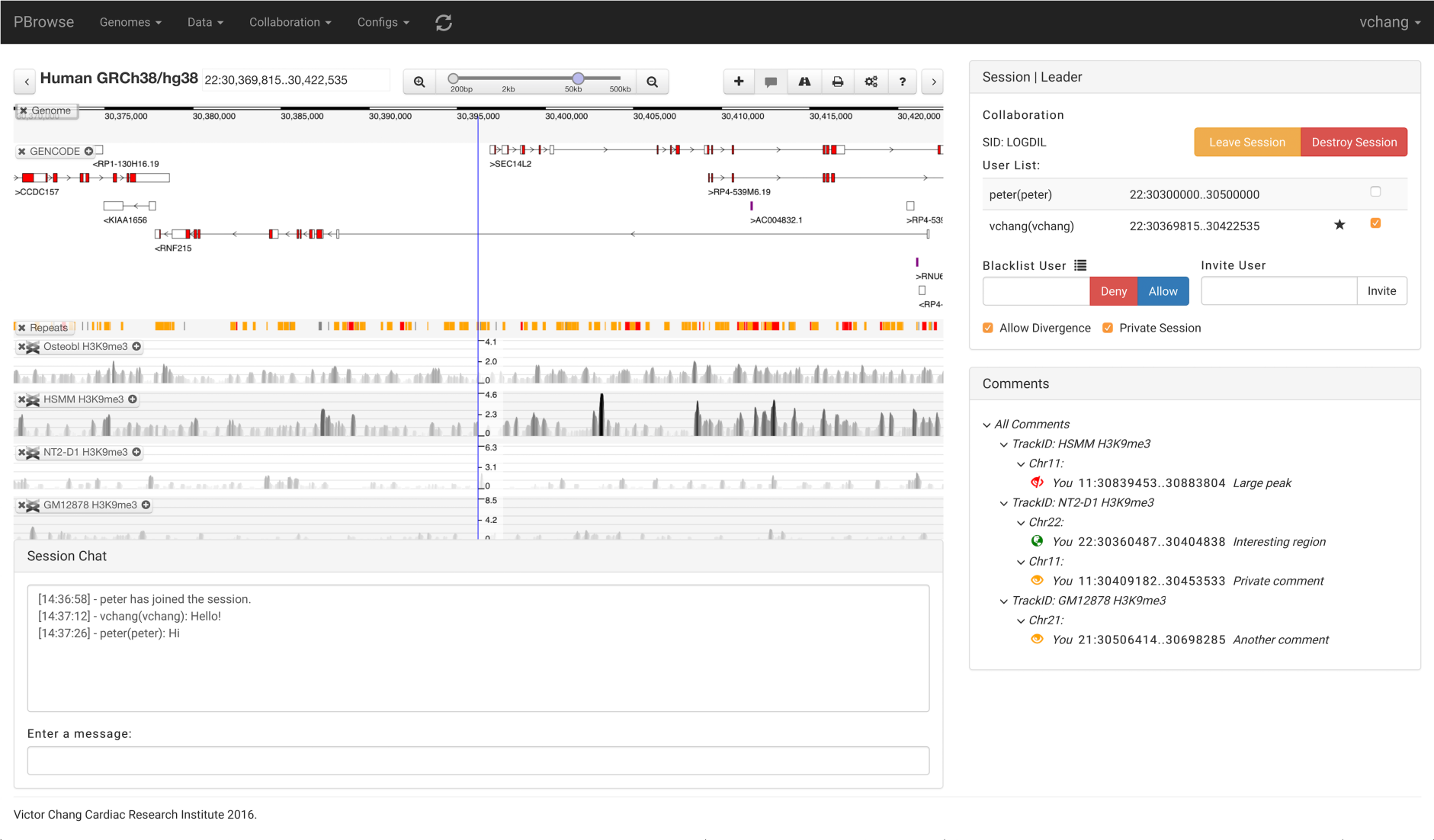
PBrowse collaborative session screenshot. Session management panel (right panel) shows information about the current collaborative session, such as the session ID, users in session and location status. Leader’s session management panel has extra functionality such as blacklisting user, inviting user and changing privacy of session. Session chat (bottom panel) allows user in the collaborative session to communicate with each other.

Individual tracks can be configured further within the browser, depending on their type, allowing for customisation of colouration, size, restrictions on the number of displayed features, among numerous other options. The genome browser stores all the track configuration information in the web-browser’s local storage, allowing past browsing sessions to be seamlessly resumed if the page is closed.

Custom tracks are also restored to their last known state, including any style changes made by the user.

### Real-time collaboration

The primary purpose of this browser is to enable real-time collaboration among multiple users, in order to improve the efficiency of data exploration tasks. This is achieved by having multiple users connected together while viewing a dataset, as part of a parallel session. All the users in the session can see the real-time status of all other users, which is updated whenever someone performs an action which modifies their view. This process is depicted in Figure 5. Participants can subscribe to the updates of zero or more users. Any updates received from a subscribed user will cause the subscriber to automatically synchronise their view to the new position. Otherwise, only the user’s record of the position is updated. The view change of the subscriber triggers a reverse update, which informs the initial user of the change to the subscriber’s position.

**Figure 5.**
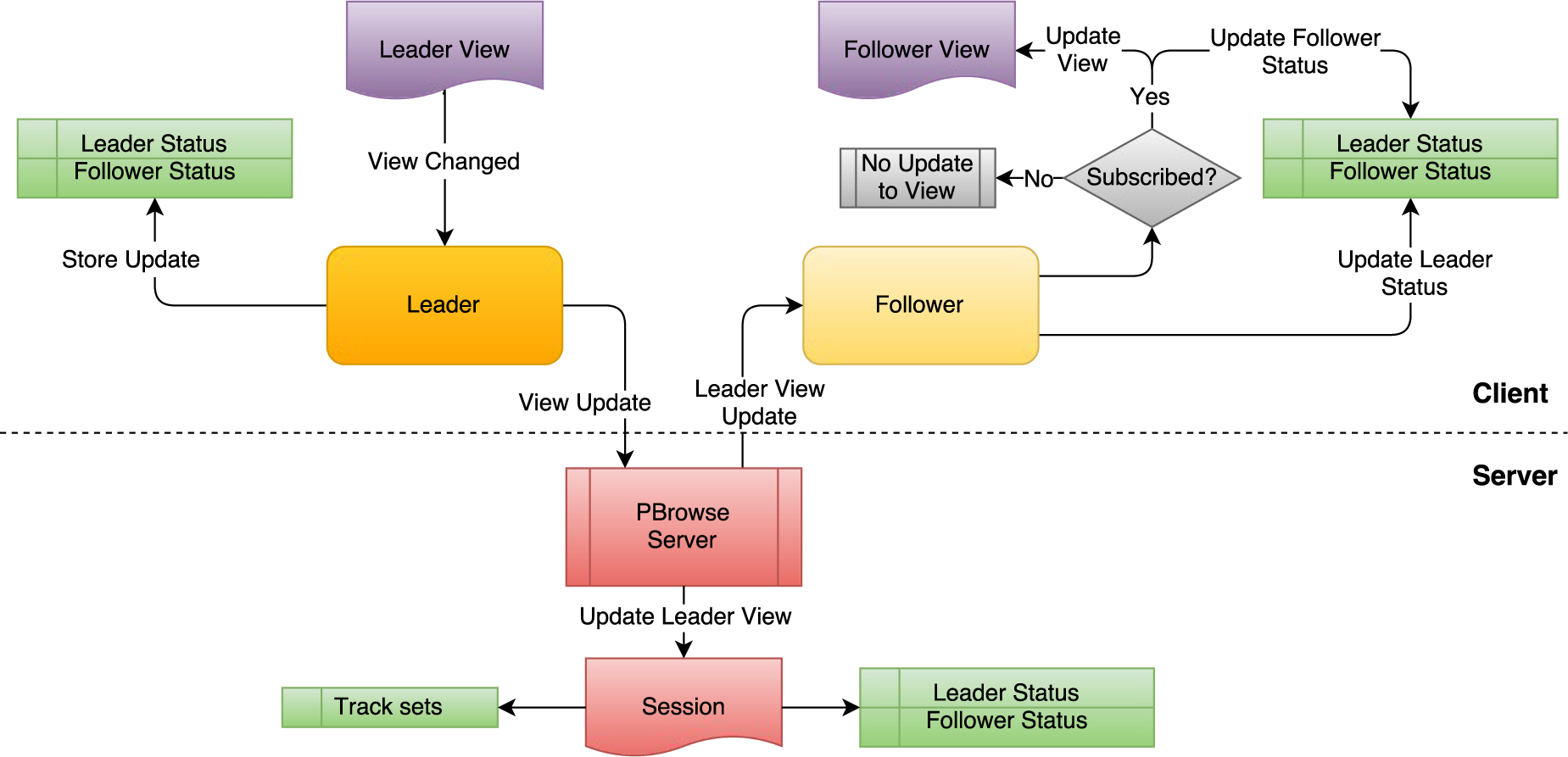
Collaborative synchronisation flowchart. Propagation of changes made by leader in their genome browser to follower’s genome browser.

When making a new parallel session, the creator is assigned the role of leader and as such, has more privileges than that of any other participant, i.e. a follower. The leader is able to perform a number of actions affecting other users in the session, including: modifying the session’s public status, inviting new users to join, or temporarily removing (kicking out) existing users, blocking other users from joining or re-joining the session, preventing follower interaction, or even terminating the session entirely. The leader may also elect their replacement from among their followers who effective immediately, becomes the new session leader – the old leader is demoted to follower status. If the current leader leaves the session at any point, the leadership position is passed onto a random follower. If the original leader returns, they will resume the position. The original leader can only be the person who created the session, or one the leader has elected to replace them. If all users leave the session, it will immediately terminate and cannot be re-joined.

By default, the session leader is not subscribed to the updates of any user although they may choose to do so. A newly connected user, however, will initially be subscribed to updates from the leader, though they may also modify their subscription set at will. Subscriptions affect only view changes; track modifications affect every user in the session equally and cannot be ignored. This covers all instances whereby the set of visible tracks in the genome browser are modified, including the reordering, removing or adding of new tracks.

The visibility of a session is controlled by its privacy status. If marked public, the session will appear as a listing available to all PBrowse users. They will see its identifier and whether or not the session requires a passcode. If it does require a code, users must enter it, otherwise, any user may join the session. Private marked sessions will not be listed, requiring a user to know both the identifier and passcode – if any. Users invited to join a session, however, need only to accept the invitation. If a user has been blacklisted from a session, they cannot join it, under any circumstance.

In addition to synchronising views, all the members of a collaborative session are granted temporary access to files uploaded by all other group members. This includes those who were part of the session but since left it. It affects all user uploaded files, except those explicitly marked “only-me” – which as the name indicates, are visible only to the uploader. This facilitates the quick sharing of new data, requiring the file to be uploaded only once before it is viewable by multiple collaborators. While participating in a collaborative session, the users may also communicate directly via the session chat feature. It allows them to send short textual messages to all other participants instantaneously. Users also receive session status updates as special messages i.e. when users join, leave, are kicked out, etc. This can be useful for communicating intent or discussing collaborative tasks.

### Track comments

The ability of users to leave comments on data tracks is another core feature of the collaborative framework of PBrowse. In this case, however, it is both real-time and deferred. When a user adds a custom track, they unlock the option to register a new, persistent comment on the currently viewed genomic region. The commenter can specify any message to accompany it, or simply leave it blank – in which case it may serve as a convenient bookmark to a specific region of interest. The commenter can also specify to make the comment public, private, or visible only to the author i.e. only-me. If public, when the comment is made, all users currently viewing the track, regardless of whether they share a collaborative session, will be notified of its existence. If private, the comment will only ever be visible to the author, or those within the same collaborative session. Finally, if the comment is marked “only-me”, it will only ever be visible to the author, and it will not be shared under any circumstance. Consequentialy, public comments made on private tracks will only be visible users with the permission to view the track, i.e. those within the same collaborative session.

Comments are displayed in a tree-like view, with a single root node having a branch for each currently loaded track. Under each track node, comments are further grouped by the chromosome under which they appear. At the lowest level, the full comment details are visible, including its author, its exact location, and any textual annotation. Clicking on any node will expand or contract it, while clicking on a comment node will open a menu allowing for further operations. All users can use this to immediately navigate to the region of interest, while only the comment author is able to delete it. When a user loads a track for the first time, all of the public comments on that track are retrieved from the database and rendered. Removing the track from view will subsequently hide its associated comment branch. Only comments made while actively viewing the track are considered new, as the read status of a comment is not persistently tracked.

### Custom tracks

The Dalliance browser alone offers some means of managing data tracks, and adding new ones from external sources, primarily via the Distributed Annotation Server (DAS) protocol. These data are served as plain XML to the client, which processes and renders it as a track within the primary view, in a form according to the fetched data type e.g. gene or sequence. Dalliance can additionally render tracks from indexed binary data files served directly from HTTP, supporting the 2bit, bigWig, bam and bigBed formats (14) with the possibility for random access via range queries. The BED, WIG, and VCF formats are also accessible in this way, but they must be loaded in their entirety into memory before particular regions can be retrieved. This is undesirable as the client must waste bandwidth transferring all non-essential parts of the file. The only current workaround to this is to convert the data into its associated binary form, i.e. WIG to bigwig, BED to bigBED, etc. or provide a tabix index along with the compressed data file. The binary conversion approach is made easily possible by a number of publically available utilities provided by UCSC and is the recommended approach when visualising such data in PBrowse.

A number of different genomes are made available for browsing within PBrowse, the default being the GRCh37/hg19 human genome, which has been preloaded with a large dataset (2,485 tracks) from the ENCODE project (15), ready for immediate viewing. In addition, PBrowse currently also provides the hg18, and GRCh38/hg38 human genomes, the Zv9 zebrafish genome, the GRCm37/mm9and GRCm38/mm10 mouse genomes, and the WS220 worm genome. New tracks can be added by any user in a process described in the following section. Newly added data is categorised among other sources which share the same reference genome.

New genomes may be added by users through a simple, two-step procedure. First a data file containing sequence information for the particular genome must be uploaded. Then the user changes the active genome to generic via the drop-down menu, and follows the prompts, providing a name, as well as the sequence file ID, and an optional gene file ID. The new genome exists temporarily but can be preserved by saving the current track configuration. The saved configurations are stored by PBrowse and recalled for each user, who can have multiple save states. They can also be shared with other users who are in the same collaborative session.

### Data upload and sharing

In addition to collaboration, PBrowse also provides an efficient means for users to upload and share custom track data. When initiating an upload, users can specify: a “track-name” – the primary means of identifying the data, its description – which can contain any extra relevant information relating to the annotation, a study ID – which allows for easy grouping of related data files, the genome tag – which states the data’s associated genome assembly e.g. hg38, and finally the file’s public status – which determines its visibility to other users of PBrowse. The genome tag acts only as a suggestion for other users to follow when visualising the data. The file metadata is tabulated and presented to the user who can sort entries based on the contents of particular columns, or perform keyword searches to locate specific tracks, as depicted in Figure 6.

**Figure 6.**
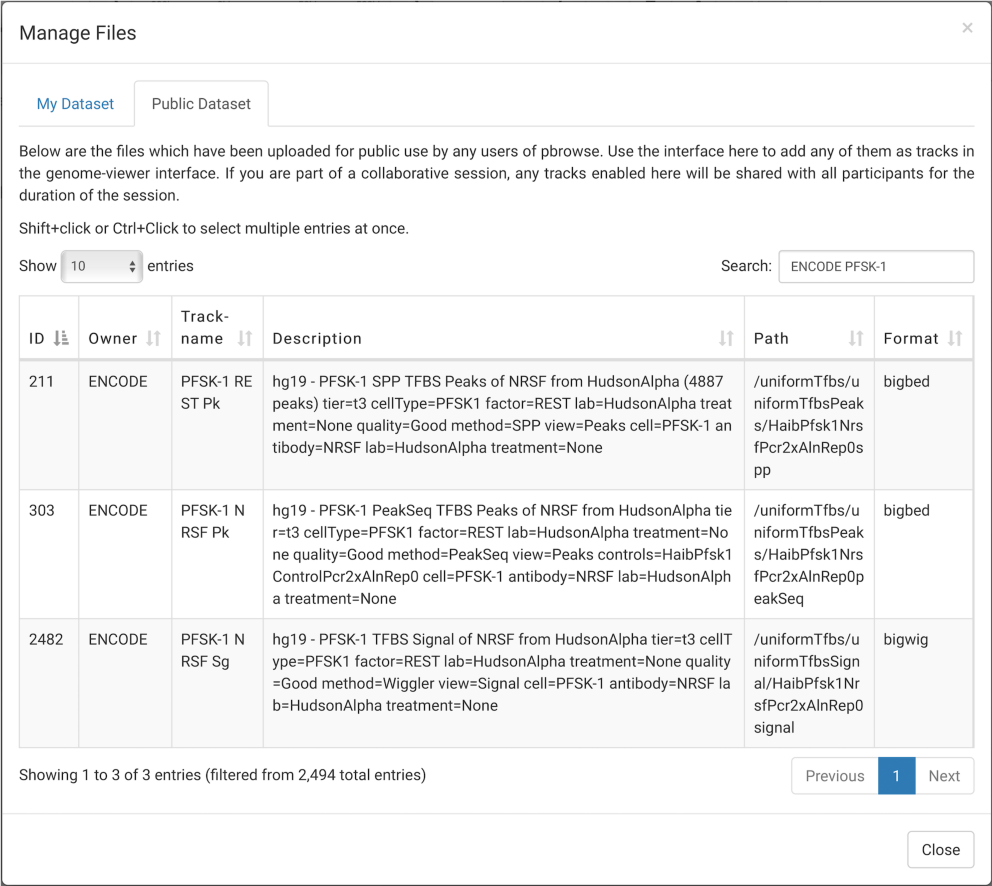
File management interface. User is able to access their uploaded files (My Dataset) and files made public by other user, such as the ENCODE dataset, (Public Dataset) and add them as tracks to PBrowse.

The visibility of an uploaded file can have one of three distinct values, “only-me”, “private” or “public”. If a file is set to only-me, only its owner, i.e. the uploader, can see it or visualise it. The same applies to the private setting, except during a collaborative session where all users share their privately accessible files automatically with other members of the session – including those uploaded during the session. This sharing remains until the user leaves the session, in which case they will lose access to files shared by the collaborators. Meanwhile, the remaining users will still be able to access the leaving user’s files until the session is completely concluded.

Publicly accessible files, on the other hand, are always visible, to every user of PBrowse, whether they are collaborating with the file owner or not. Users do not even need to be logged in to visualise them, although most functionality will be restricted without doing so.

The level of access granted to collaboratively shared files is the same as the access granted on publically visible files. In both cases, the data can only be added to the genome browser as a track. However, when managing their own uploaded files, a user is additionally able to modify the privacy status, share it with a specific group, or delete the file entirely. Any changes affecting a file’s visibility are propagated in real-time, with all affected users being notified as well.

As an alternative to uploading files to the PBrowse server, it is possible to simply register the location of a remotely-hosted file. The user can still provide meta-information for the source and a corresponding file listing will be produced and displayed within the file manager. All standard file management options can be applied to a remotely registered file, including privacy settings and search filtering, with the obvious exception that deleting the entry will not destroy the remote source. Once uploaded or registered, clicking on a listing within the file manager will present the user with the available options for the file. Multiple files can also be selected at once to perform batch operations. However, this requires the user to have the correct privilege to perform the action for all selected files, otherwise, the operation will fail. The batch management feature is particularly useful for adding multiple related tracks to the browser at once.

## 4 CASE STUDY: COLLABORATIVE EXPLORATION OF HUMAN CARDIAC ENHANCERS WITH ENCODE DATA

To demonstrate how PBrowse enables collaborative exploration of genome-wide data, we constructed a case study involving sharing and exploration of EP300 ChIP-seq data set from foetal and adult human heart tissues (15). We have created a short video of this demonstration and it is available in Supplementary material S1.

In the case study, we have a user (Xin) who has loaded a few EP300 tracks from human and foetal heart in his genome browser. The user then starts a new collaborative session and invites his collaborator (Andrian) to join the session. Upon joining the collaborative session, the collaborator’s genome browser view is synchronised to the view of the user genome browser, allowing the collaborator to view the same EP300 tracks at the same view. Changes made by the user on his genome browser, such as scrolling to a different region and an addition of the H3K27ac tracks from the user’s computer and ENCODE DAS server, are reflected on the collaborator genome browser instantly demonstrating the real-time nature of the collaborative session. Finally, the case study also demonstrates the other collaborative features of PBrowse, such as instant messaging – allowing the user to communicate with the collaborator in real-time – and comments – which allows for users to share the region of interest with their collaborators and vice versa.

## 5 CONCLUSION

The technology of web-based genome browser has evolved significantly over the last decade. Nonetheless, the focus of these browsers has been largely around the personal exploration of data. To fully harness the collective wisdom of a collaborative group, PBrowse places a strong emphasis on human-human interactions through the medium of the genome browser. As illustrated in our case study, PBrowse enables a level of real-time knowledge exchange between multiple users that are not currently achievable by other browsers. Thus, we argue that PBrowse represents a paradigm shift in the way we see and use genome browser. A public demonstration version is available at http://pbrowse.victorchang.edu.au, while the source code is available at GitHub, http://github.com/VCCRI/PBrowse, under BSD 3 license.

## ACKNOWLEDGEMENT

We thank the IT department of the Victor Chang Cardiac Research Institute for providing technical support. We also thank the Information and Communication Technology Office (ICTO) at the University of Macau for providing access to and support on a High-Performance Computer, and Jacky Chan and William Pang for their expert technical support.

